# CNTNAP2 is targeted to endosomes by the polarity protein Par3

**DOI:** 10.1101/601575

**Authors:** Ruoqi Gao, Christopher P. Pratt, Sehyoun Yoon, Maria Dolores Martin-de-Saavedra, Marc P. Forrest, Peter Penzes

## Abstract

A decade of genetic studies has established Contactin-associated protein-like 2 (*CNTNAP2*) as a prominent susceptibility gene associated with multiple neurodevelopmental disorders. The development and characterization of Cntnap2 knockout models in multiple species have bolstered this claim by establishing clear connections with certain endophenotypes. Despite these remarkable in vivo findings, CNTNAP2’s molecular functions are relatively unexplored, highlighting the need to identify novel protein partners. Here, we characterized an interaction between CNTNAP2 and Partitioning-defective 3 (Par3) – a polarity molecule we isolated in a yeast-two hybrid screen with CNTNAP2’s C-terminus. We provide evidence that the two proteins interact via PDZ domain-mediated binding, that CNTNAP2^+^/Par3^+^ complexes are largely associated with clathrin-coated endocytic vesicles, and that Par3 causes an enlargement of these structures. Live imaging and fluorescence recovery after photobleaching (FRAP) reveals that Par3 limits the mobility of CNTNAP2 at endosomes, thus stabilizing it at that location. Finally, expression of Par3 but not Par3ΔPDZ can cluster endogenous CNTNAP2 in primary neurons. Collectively, we conclude that Par3 regulates CNTNAP2 spatial localization to endocytic compartments.

## 1. Introduction

*CNTNAP2* is a prominent disease susceptibility gene associated with Autism Spectrum Disorder (ASD), Intellectual Disability (ID), Schizophrenia (SCZ), and epilepsy, and is causative for Cortical Dysplasia Focal Epilepsy (CDFE) syndrome (Rodenas-Cuadrado et al., 2014). Similarly, *Cntnap2* knockout mice, zebrafish, and rats have recapitulated several disease endophenotypes, including impaired social interaction, communication, and seizures (Penagarikano et al., 2011; Hoffman et al., 2016; Thomas et al., 2017). *CNTNAP2* encodes a type I transmembrane cell adhesion molecule that has a wide range of functions, including clustering potassium channels, migration, and dendritic stabilization (Rodenas-Cuadrado et al., 2014). Moreover, multiple reports have shown that CNTNAP2, as well as close family members, are more prominently expressed in interneurons and preferentially affect interneuronal function (Mo et al., 2015; Karayannis et al., 2014; Gao et al., 2018). Indeed, interneuron dysfunction may be key factor for behavioral endophenotypes in *Cntnap2* null mice (Jurgensen and Castillo, 2015; Bridi et al., 2017; Vogt et al., 2014); however, the cellular and molecular mechanisms remain poorly understood.

Polarity is an organizing process that is essential for cellular development, allowing cells to form asymmetrical specializations critical for cellular differentiation (Goldstein and Macara, 2007). The partitioning defective (Par) proteins participate in an evolutionarily-conserved complex which initiates many of these essential processes through the interaction of its three members – Partition-defective protein 3 (Par3, a.k.a. PARD3), Partitioning-defective 6 (Par6), and atypical protein kinase C (aPKC) – and downstream guanine nucleotide exchange factors (Johansson et al., 2000; Lin et al., 2000). Through this interplay, Par3 assists in regulation of multiple polarity events including apical-basal establishment in epithelial cells, axon specification in neurons, asymmetric mitotic spindle positioning during cell division, and astrocyte migration (Mertens et al., 2006; Macara, 2004). Conversely, Par3 dysfunction leads to loss of polar organization, abnormal GTPase signaling, and carcinogenesis/metastasis in some cell types (McCaffrey et al., 2012).

Other than its canonical role with Par6 and aPKC, Par3 also has Par complex-independent functions. For example, Par3 can spatially localize the Rac-GEF TIAM1 to the membrane, a process important for events requiring cytoskeletal rearrangement, such as spinogenesis (Duman et al., 2013; Zhang and Macara, 2006), migration (Narayanan et al., 2013), and tight junction formation (Chen and Macara, 2005; Narayanan et al., 2013). Par3 also is involved in mammalian cell survival and phosphoinositide signaling through membranous interactions with exocyst docking proteins (Ahmed and Macara, 2017) and PTEN respectively (Feng et al., 2008). Previous studies have also implicated cell adhesion molecules, such as JAMs, Ephrin receptors, and GPCRs, as critical upstream effectors for recruitment of Par3 (Itoh et al., 2001; Lin et al., 1999; Duman et al., 2013). Thus, Par3, in both canonical and non-canonical roles, serves as a central recruitment scaffold for a multitude of interactors, many of which likely remain unknown.

Here, using a yeast two-hybrid screen of CNTNAP2’s C-terminus, we discovered Par3 is a novel CNTNAP2 interaction candidate. We validated this interaction by co-immunoprecipitating Cntnap2 and Par3 from mouse brain. Through domain mapping in HEK293T cells, we demonstrated this association to be dependent on PDZ binding. Immunocytochemical analysis in COS7 cells revealed both molecules in association within abnormally large clathrin-dependent endocytic vesicles, while live imaging showed that this association altered CNTNAP2 mobility. Finally, overexpressed Par3 formed abnormally large vesicular structures in primary cultured neurons, some of which were enriched in endogenous CNTNAP2. Taken together, we characterized Par3 as a novel CNTNAP2 interaction partner, which may contribute to a deeper fundamental understanding of CNTNAP2’s molecular functions and associated biological pathways that contribute to neurodevelopmental disorders.

## 2. Materials and Methods

### Antibodies and Plasmids

A detailed antibody and primer list is presented in the Supplementary Information.

pEGFP-N2 plasmid was purchased from Clontech (Mountain View, CA, USA). pEBFP2-N1 was purchased from Addgene (Cambridge, MA, USA; #54595). FLAG-CNTNAP2 was generated by subcloning human CNTNAP2 cDNA (gift from Dr. Elior Peles, Weizmann Institute of Science, Israel) into the pEGFP-N2 vector (restriction sites BamHI and NotI, removing EGFP) using the Infusion cloning system from Clontech. The FLAG sequence was then inserted downstream of the signaling peptide (amino acids 1-27) using the same technique. mCherry-CNTNAP2 was constructed similarly, with mCherry being inserted after the signal peptide. CNTNAP2 truncations were created by deleting the C-terminus, 4.1-binding site, or PDZ-binding site.

Myc-Par3 was created by subcloning human myc-Par3 from the pK-myc-Par3b plasmid (Addgene #19388) into the peGFP-N2 vector as described above using BamHI and NotI restriction sites. Par3 truncation mutants were created by deleting the first (amino acids 271-359), second (amino acids 461-546), third (amino acids 590-677), or all PDZ domains (ΔPDZ_all_; amino acids 271-677). GFP-Par3 and GFP-Par3ΔPDZ_all_ were generated by inserting EGFP into myc-Par3 between the BglII and BamHI restriction sites.

### Neuronal Culture and Transfections

High density (300,000 cells/cm2) cortical neuron cultures were prepared from Sprague-Dawley rat E18 embryos. Cortical neurons were transfected using Lipofectamine 2000 (Thermo Fisher Scientific, Waltham, MA, USA) following the manufacturer’s recommendations. Neurons were maintained in feeding media for 3 days post-transfection. Any visual signs of poor neuronal health meant the exclusion of the cell from quantification. COS7 or HEK293T cells (ATCC, Manassas, VA, USA; lines authenticated before shipment) were transfected using Lipofectamine 2000. Experiments were performed 48h post-transfection.

### Immunocytochemistry

Endogenous Par3 was detected as described (Zhang and Macara, 2006). Briefly, neurons were fixed in 4% formaldehyde-sucrose-PBS for 15 min at room temperature (RT), followed by permeabilization with 0.2% Triton-X-100 for 5 minutes at RT, blocking in 10% bovine serum albumin (BSA) for 1h at RT, and incubation of primary antibody in 3% BSA overnight at 4°C. Secondary antibodies were applied in 3% BSA 1h at RT. For detection of all other targets, ICC was performed as described (Gao et al., 2018). Coverslips were mounted using ProLong Gold.

### Immunoprecipitation

Mouse cortex or HEK293T cells were homogenized in immunoprecipitation buffer (50 mM Tris pH 7.4, 150 mM NaCl, 0.5% Triton X-100), with protease inhibitor cocktail (Roche, Basel, Switzerland) and solubilized for 1h at 4°C. Solubilized material was centrifuged at 20,000 g for 10 minutes at 4°C and the supernatant was precleared with protein A/G sepharose beads (Thermo Fisher Scientific) for 30 minutes. Proteins were then immunoprecipitated with 3 *μ*g of antibody overnight at 4°C, followed by a 1 hour incubation with protein A/G sepharose beads the following day. Beads were then washed 3 times with IP buffer before adding 2x Laemmli buffer (Biorad, Hercules, CA, USA). Samples were analyzed by SDS-PAGE and Western blotting.

### Fractionation

Subcellular fractionation was performed as previously described (Nakagawa et al. 2005). Briefly, cortices from 6-week old mice were homogenized in cold sucrose buffer (20 mM HEPES pH 7.4, 320 mM sucrose, 5 mM EDTA) with protease inhibitor cocktail (Roche). Homogenates were centrifuged at 3,000 g for 20 min at 4°C speed to pellet nuclei. Supernatant (S1) was then centrifuged at 38,000 g for 30 min at 4°C to obtain a crude membrane pellet (P2). P2 was re-suspended in potassium iodide buffer (20 mM HEPES pH 7.4, 1 M KI, 5 mM EDTA) to remove membrane-associated proteins (S3). Membranes were again collected by centrifugation (38,000 g for 20 min at 4°C). Membranes were washed (20 mM HEPES pH 7.4, 5 mM EDTA) and pelleted once more (S4) before solubilizing in CHAPS buffer (20 mM HEPES pH 7.4, 100 mM NaCl, 5 mM EDTA, 1% CHAPS) supplemented with protease inhibitors for 2 hours at 4°C. Solubilized membranes were clarified by centrifugation at 100,000 g for 30 min at 4°C (S5). The final CHAPS-insoluble pellet was re-suspended in SDS buffer (50mM TRIS pH 7.4, 150 mM NaCl, 1% SDS) supplemented with protease inhibitors, solubilized at 37°C for 20 mins and clarified by centrifugation (S6).

### Yeast Two-Hybrid Screening

Y2H screening was performed using the Matchmaker Gold System (Clontech). Briefly, the C-terminal region of CNTNAP2 (amino acids 1284-1331) was cloned into the bait vector (pGBKT7) and transformed into yeast. Bait-positive yeast were then mated with yeast containing cDNA (pGADT7) from a mouse brain cDNA library. Mated yeast were then plated onto double amino acid (-Leu/-Trp) dropout, Aureobasidin A positive (A), and X-*α*-Gal positive (X) plates (DDO/X/A), a low stringency medium that selects for mated yeast containing both bait and prey vectors. DDO/X/A colonies that were blue, indicative of a possible interaction between bait-prey, were streaked onto quadruple amino acid (-Leu/-Trp/-His/-Ade) dropout, Aureobasidin-A positive (A), and X-*α*-Gal positive (X) plates (QDO/X/A), which require a genuine bait-prey interaction for growth. The blue colonies from QDO/X/A plates were isolated and plasmids sequenced.

### Confocal Microscopy

Confocal images of neurons were acquired using a Nikon C2 confocal microscope using a 60X oil immersion objective with numerical aperture (NA) = 1.4 with 0.4 *μ*m z-stacks. All images were acquired in the linear range of fluorescence intensity and a single plane image from the stack was selected for analysis. For antibody validation, z-projections were used. The acquisition parameters were kept the same for all conditions and images were analyzed using Fiji software.

### Colocalization analysis

Colocalization highlighter images and Manders’ colocalization coefficients were determined in ImageJ after thresholding. Total immunofluorescence intensity of region of interest (ROI) was measured automatically. For the size of endosome quantification, a threshold was applied to the maximum projection images to include all detectable endosomes. The endosome restricted to objects with areas greater than 0.08 *μ*m2, were manually detected, and the endosome area was measured. For object-based binary classification of colocalization, see Trafficking Marker Analysis.

### Trafficking Marker Analysis

COS7 cells were transfected with mCherry-CNTNAP2, Myc-Par3, and Blue Fluorescent Protein (BFP). Cells were fixed with standard immunocytochemical methods (see Immunocytochemistry section) and stained with respective antibodies (see Antibodies and Plasmids). Images were acquired on a Nikon C2 scanning confocal microscope using a 60X objective (NA = 1.4) at Nyquist sampling frequency. Cells were imaged after determining suitable expression of Par3, CNTNAP2, and BFP.

ROIs were drawn on single planes encompassing whole cell cytoplasm and excluding nuclei using BFP cell-fill as a guide. Images were processed and analyzed using Python 3.6 with the scikit-image package (van der Walt et al., 2014). Briefly, images were preprocessed using a 3×3 median filter and 20-pixel disk white tophat transformation to reduce noise and background, respectively. Thresholding was performed using the Triangle algorithm(Zack et al. 1977) for each color channel and converted to binary images. Small objects (¡12 pixels2) were excluded from analysis. Puncta were segmented into objects using a Watershed algorithm. Colocalization analysis was performed as follows: For two-channel colocalization of Par3 and CNTNAP2, an object in the first channel was considered colocalized with the second channel if >50% of its included pixels were also above threshold in channel 2. For triple colocalization of CNTNAP2^+^/Par3^+^ puncta with trafficking markers, the intersection of these channels was then segmented as described above. These objects were then used as channel 1 for colocalization with markers (channel 2). See Supplemental Figure 2 for illustration.

### Code Availability

Python code for trafficking marker analysis is available upon request.

### Live Imaging and Analysis

COS7 cells for FRAP experiments were grown on glass bottom culture dishes from MatTek (Ashland, MA, USA) and transfected with mCherry-CNTNAP2 and GFP-Par3 or GFP-Par3ΔPDZ_all_. Cells were imaged with a confocal microscope (Nikon) in a CO2 incubator stage. Images were taken with 256×256 pixel resolution every 1s for 300s after bleaching. 100% laser power pulses of 1ms for 1 min were used to bleach GFP-Par3. Fluorescence intensity of bleached GFP-Par3 was measured and normalized to the fluorescence before bleaching with the same area. Recovery data points were then fitted to a one-phase association exponential in GraphPad Prism (Version 7).

### Blinding and Statistical Analysis

Data from cell line and primary neuron studies were obtained and analyzed under blinded conditions (coverslip identity hidden, cells selected randomly, filenames randomized). Cells of visibly poor health were excluded from quantification. Sample sizes for cell line studies were between 10-30 cells. For primary neuronal studies, over 30-40 neurons were imaged and the best representative neuron was used for display.

All statistical tests were performed with GraphPad Prism (Version 7) or in Python using the SciPy library. Before analysis, data were first tested using D’Agostino’s Omnibus Normality Test and Pearson correlation in order to determine whether parametric or nonparametric tests were to be used. Post-hoc tests were always used in multiple comparison analysis. P values < 0.05 were considered significant.

## 3. Results

### 3.1. A yeast two-hybrid screen reveals Par3 as a novel CNTNAP2 interaction partner

We performed a yeast two-hybrid screen of a mouse brain cDNA library with the entire CNTNAP2 intracellular domain (ICD) as bait (amino acids 1284-1331) to find intracellular interactors of CNTNAP2 (Figure 1a). Out of three million candidates screened, we identified 41 unique clones (Gao et al., 2018), with Par3 being one of the most intriguing, given its role as a spatial organizer (Zhang and Macara, 2006; Duman et al., 2013; Ahmed and Macara, 2017). We confirmed the this positive result by demonstrating yeast growth on high stringency plates only when expressing both bait/prey plasmids, indicating a direct physical interaction (data not shown). We then validated that CNTNAP2 and Par3 are in the same protein complex in vivo by co-immunoprecipitation experiments from mouse cortex homogenates (Figure 1b). To determine the protein domains involved in their interaction, we co-expressed various CNTNAP2 truncation mutants (Figure 1a; red lines) with Par3 in HEK293T cells and performed co-immunoprecipitations. We found that the C-terminal PDZ-binding motif of CNTNAP2 is necessary for interaction with Par3, as its deletion prevented Par3 pull-down (Figure 1c). Conversely, deletion of any one or all of Par3’s three PDZ domains (Figure 1a; red lines) resulted in a dramatic reduction of CNTNAP2 co-immunoprecipitation efficiency (Figure 1d). From these results, we conclude that CNTNAP2’s C-terminal PDZ-binding domain can interact with multiple Par3 PDZ domains, a Par3 binding motif previously observed with Par6 (Renschler et al., 2018). We decided to use a Par3 mutant lacking all PDZ sites (Par3ΔPDZ_all_) for subsequent studies. These experiments demonstrate that Par3 is a novel CNTNAP2 interactor that utilizes PDZ binding for physical association.

**Figure 1:**
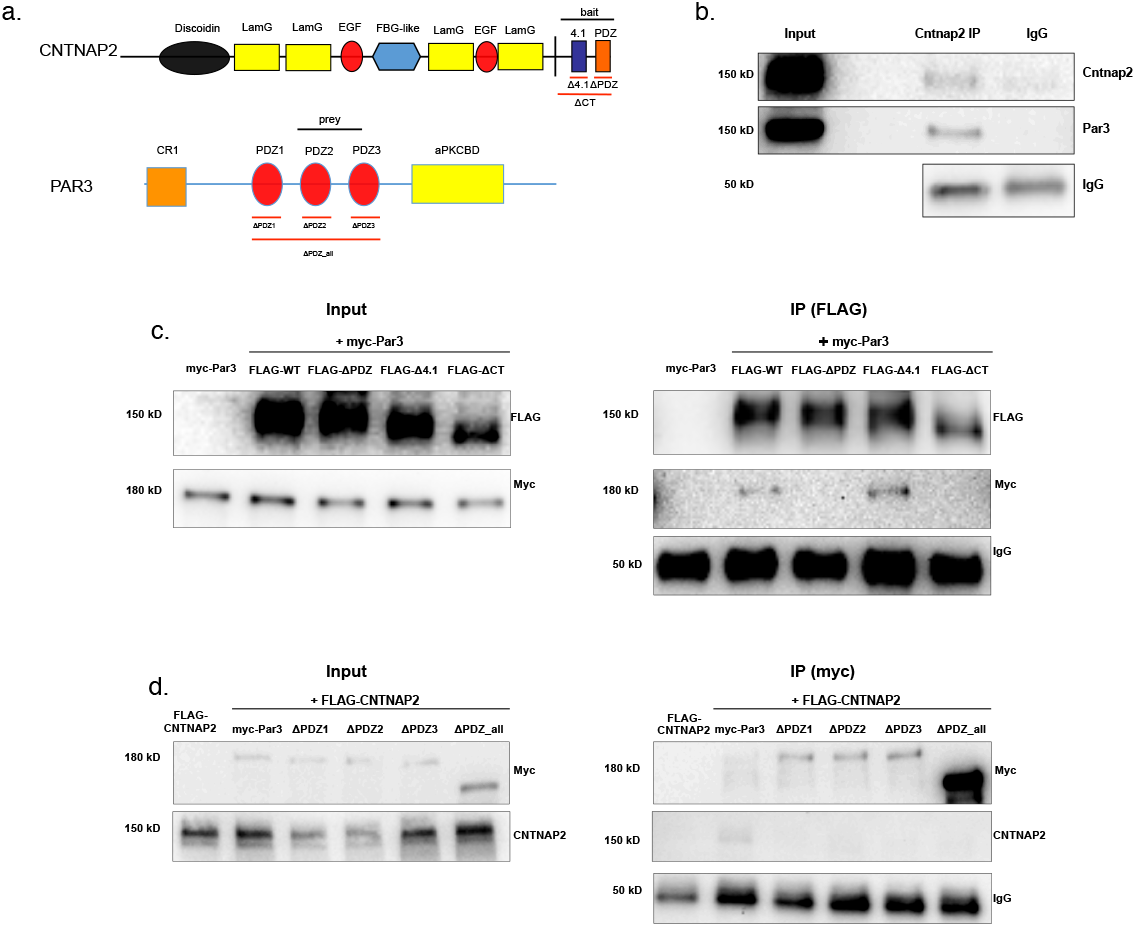
Par3 binds with CNTNAP2 through PDZ interaction. (a) Cartoon showing the domain topology of CNTNAP2 and Par3, with black lines representing the bait/prey regions derived from the yeast-two hybrid screen and red lines indicating the truncated mutants used for experimentation. (b) Representative western blot showing Par3 co-immunoprecipitation with CNTNAP2 from mouse cortex homogenates (representative image from 2 independent experiments). (c) Western blot showing co-immunoprecipitation of Par3 coexpressed with various CNTNAP2 truncations to demonstrate this interaction is specific to CNTNAP2’s PDZ-binding domain (representative image from 3 independent experiments) (d) Western blot showing co-immunoprecipitation of CNTNAP2 with various Par3 truncations to demonstrate each of Par3’s three PDZ domains can bind to CNTNAP2 (representative image from 2 independent experiments).

### 3.2. Subcellular compartmentalization of Par3 and CNTNAP2

We proceeded to characterize the spatial properties of both proteins by performing an in-depth subcellular fractionation procedure using salt extraction and detergent solubilization (Figure 2). To confirm fractionated compartments, we probed the fractions with GluA1, PSD95, and beta-tubulin antibodies. GluA1 (synaptic membrane protein) was highly enriched in the crude membrane (S5) and synaptic (S6) fractions while PSD95 (synaptic cytosolic protein) was only enriched in the S6 fraction; beta-tubulin (non-synaptic cytosolic protein) was absent from either S5 or S6 fractions (Figure 2b, bottom). We found both Par3 and CNTNAP2 in the S6 fraction, consistent with previous publications (Chen et al., 2015; Oiso et al., 2009; Gao et al., 2018; Lin et al., 2000). Moreover, Par3 was also highly enriched in the crude membrane-associated (S3) fraction but almost nonexistent in S5, while Cntnap2 was most abundant in the S5 fraction, as expected. Together, these data suggests Par3 is a cytosolic protein, which can transiently associate with membrane proteins like CNTNAP2.

**Figure 2:**
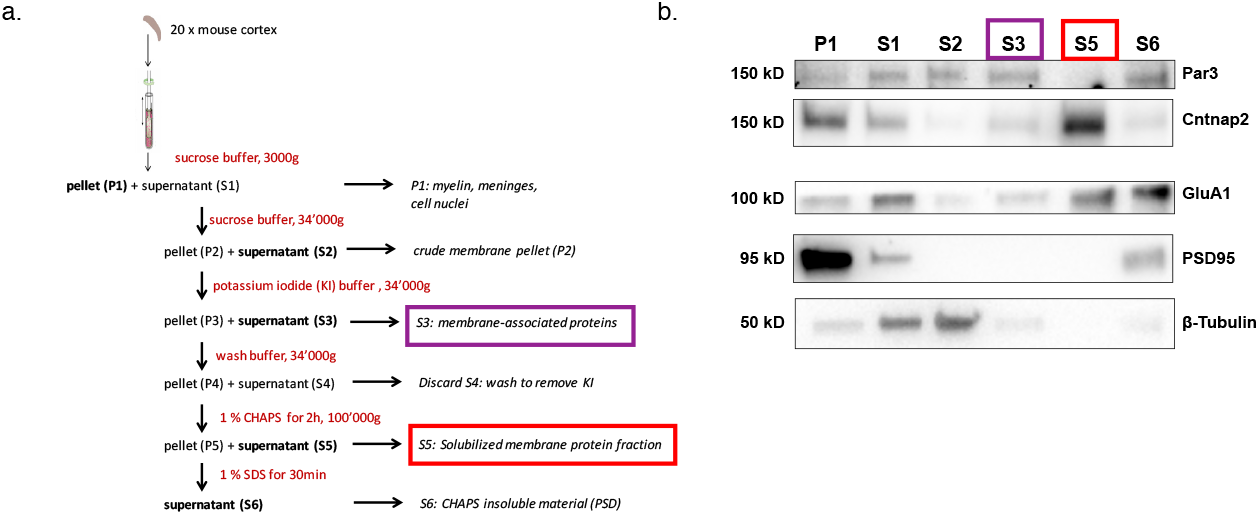
CNTNAP2 and Par3 are localized to membranes in mouse brain. (a) Schematic showing the subcellular fractionation protocol. (b) Immunoblots of subcellular fractionations from adult mouse forebrain probed with CNTNAP2 and Par3. Purple and red boxes highlight fractions of interest. GluA1, PSD95, and *β*-tubulin served as compartment-specific markers.

### 3.3. Par3 redistributes CNTNAP2 in COS7 cells

We next used COS7 cells to ascertain localization patterns for both molecules. When CNTNAP2 was expressed alone, we noted distribution patterns at the cell periphery, perinuclear region, and within small cytoplasmic vesicles (Figure 3a; mCherry-CNTNAP2 alone). Par3, on the other hand, displayed mostly a diffuse cytoplasmic localization pattern (Figure 3a; myc-Par3 alone) (Lin et al., 2000). Upon co-expression, respective expression patterns changed drastically, where both molecules were found mostly within large punctate cytoplasmic structures (arrows, Figure 3a; mCherry-CNTNAP2 + myc-Par3). Moreover, CNTNAP2 puncta size increased significantly upon Par3 co-expression (mCherry-CNTNAP2 + myc-Par3: 1.16 ± 0.04*μ*m^2^ vs. mCherry-CNTNAP2 alone: 0.46 ± 0.014*μ*m^2^; Figure 3d). These patterns were abolished with replacement of CNTNAP2 or Par3 with CNTNAP2ΔPDZ or Par3ΔPDZ_all_ mutants, respectively (Figure 3a; mCherry-CNTNAP2ΔPDZ + myc-Par3 and mCherry-CNTNAP2 + myc-Par3ΔPDZ_all_). Quantitative Manders colocalization analysis confirmed these observations (Figure 3b-c; mCherry-CNTNAP2 + myc-Par3 M2: 0.921 ± 0.02 vs. mCherry-CNTNAP2 + myc-Par3ΔPDZ_all_ M2: 0.176 ± 0.03). Our data therefore suggests a possible role of Par3 in regulating CNTNAP2 subcellular localization.

**Figure 3:**
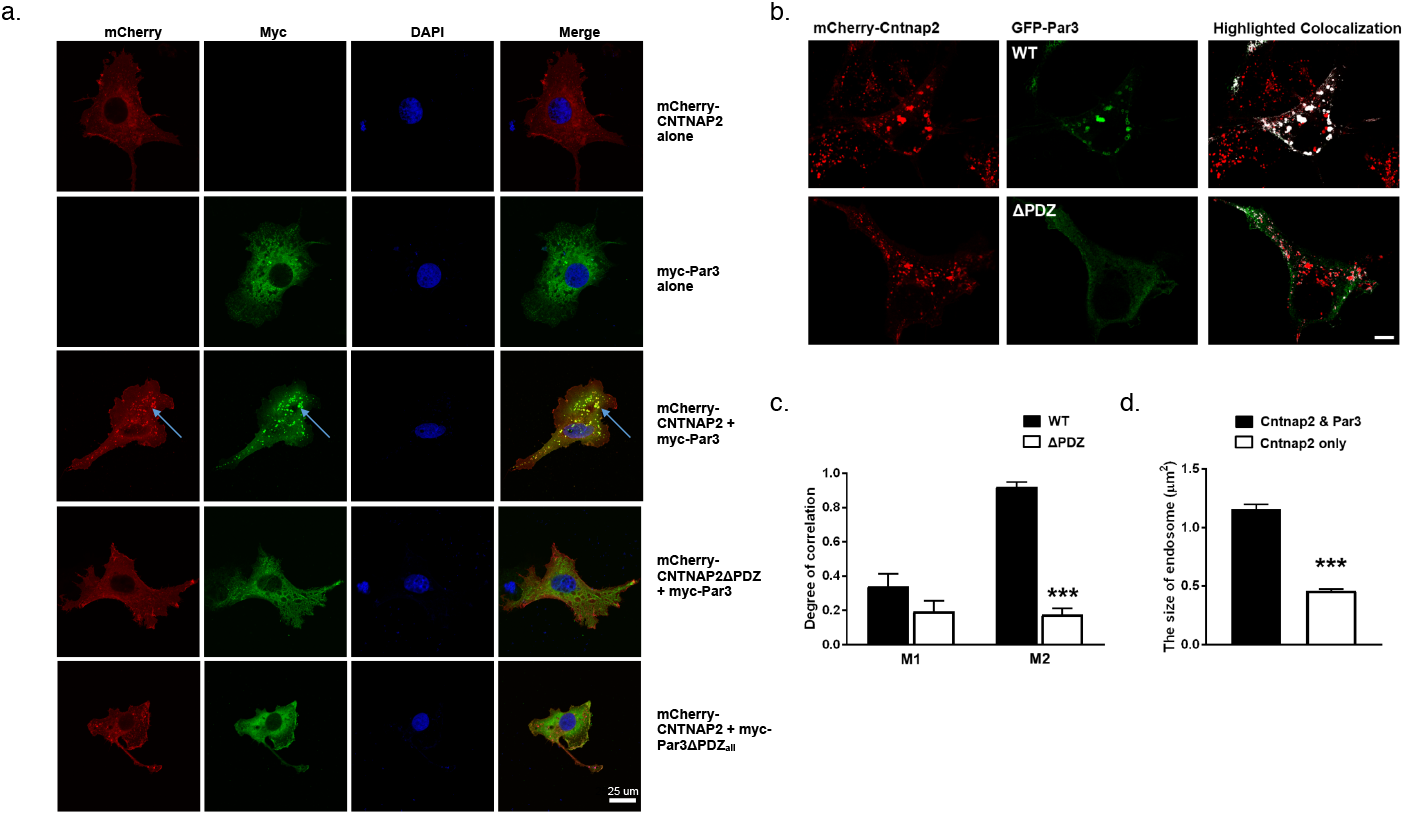
Co-expressed CNTNAP2 and Par3 associate with large vesicles in COS7 cells. (a) Representative confocal images of COS7 cells expressing mCherry-CNTNAP2/myc-Par3 alone, together, or with various truncation mutants shown to affect protein-protein interactions. Arrows point to the abnormally large mCherry-CNTNAP2 + myc-Par3 aggregates induced by co-expression of the two (scale bar = 25 *μ*m) (representative image from 2 independent experiments). (b) Selective representative images showing co-localization (white) between mCherry-CNTNAP2 + GFP-Par3 (WT) or mCherry-CNTNAP2 GFP-Par3ΔPDZ_all_ (ΔPDZ) (scale bar = 10 *μ*m). (c) Quantification of Mander’s colocalization (WT: 9 cells, ΔPDZ: 10 cells from 1 independent experiment) by unpaired two-tailed t-test (d) Measurement of the size of mCherry-CNTNAP2 vesicles alone or when GFP-Par3 is co-expressed (n = 9 cells for both conditions from 1 independent experiment) by unpaired two-tailed t-test. Values are means ± SEM. *** *P* < 0.001.

### 3.4. CNTNAP2 and Par3 co-associate with clathrin-mediated endocytic vesicles

Because the structures containing both CNTNAP2 and Par3 resembled trafficking compartments, we next determined the identity of CNTNAP2^+^/Par3^+^ structures by co-staining exogenously-expressed proteins in COS7 cells with endogenous trafficking markers. Markers for endocytosis (Caveolin-1, Clathrin heavy chain [CHC]), very early endosomes (APPL1), early endosomes (EEA1, Rab5), and the Golgi-secretory complex (GOPC, Syntaxin-6 [Stx6]) were examined. Association of CNTNAP2^+^/Par3^+^ puncta with these markers was measured using a novel object-based algorithm (Supplementary Figure 1a, also see Materials and Methods). We found CNTNAP2^+^/Par3^+^ vesicles to have a high degree of co-localization with EEA1, Rab5, and CHC (Figure 4a); line scans and quantitative co-localization analysis confirm synchronous peaks of fluorescence and statistically significant overlap between all CNTNAP2+/Par3+ and each of these three markers (Figure 4b-d). On the other hand, the remaining markers showed divergent fluorescent intensity peaks and low overlap values (Supplementary Figure 1b-c). Collectively, our analysis shows CNTNAP2^+^/Par3^+^ vesicles may be involved in clathrin-mediated endocytosis and subsequent endosomal trafficking.

**Figure 4:**
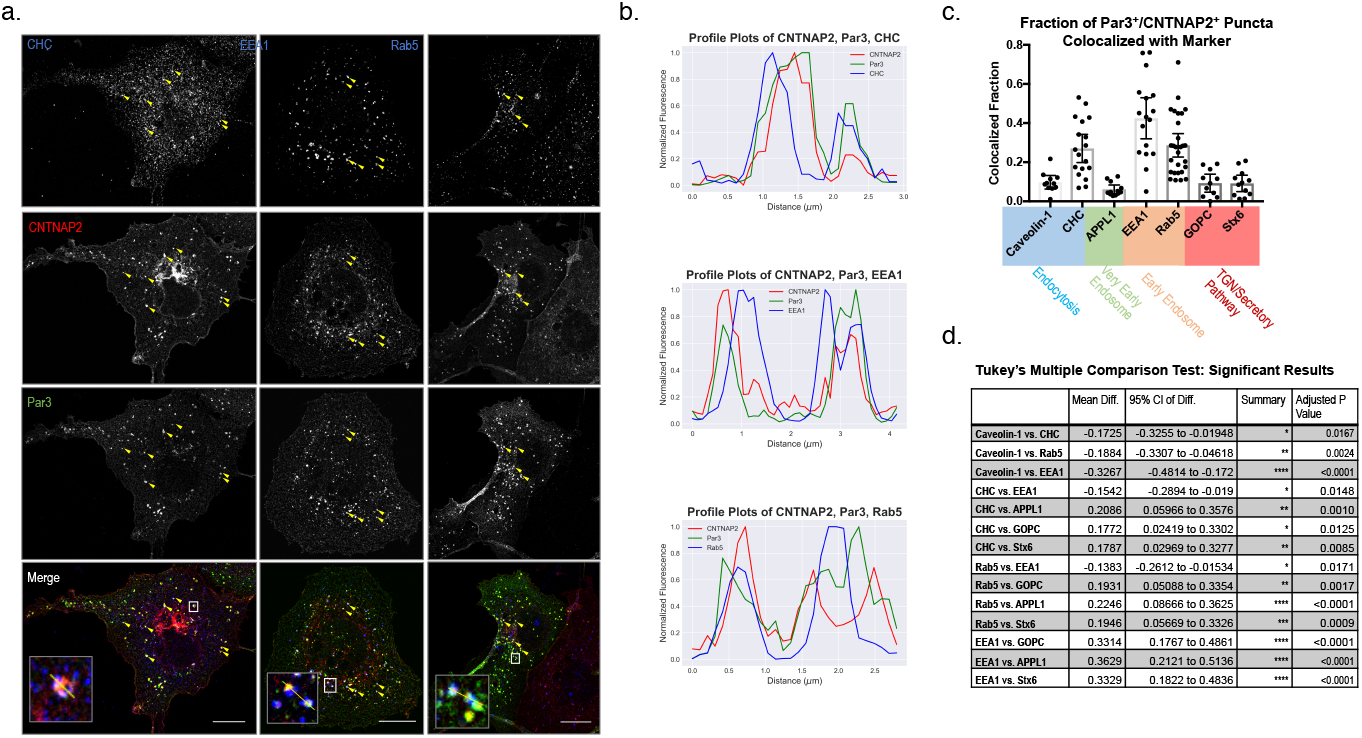
CNTNAP2 and Par3 co-localize in clathrin-coated endocytic vesicles. (a) Representative images of COS7 cells transfected with mCherry-CNTNAP2 (red channel) and Myc-Par3 (green channel) and stained with the indicated markers (far red channel). Colocalized puncta are shown with yellow arrowheads and white boxes (Scale bars =15 *μ*m) Insets at bottom are magnified images of the white boxes with line scans shown in yellow. (b) Profile plots from line-scans in (a) showing overlap of CNTNAP2, Par3, and markers. Profiles are normalized to maximum fluorescence. (c) The fraction of thresholded CNTNAP2^+^/Par3^+^ puncta showing colocalization with marker (see methods and Supplementary Figure 1). Color-coding below indicated relevant intracellular compartments. (d) Summary table showing all significant differences in means by Tukey’s Multiple Comparison Test (Caveolin 1: 11 cells, CHC: 18 cells, Rab5: 28 cells, EEA1: 17 cells, GOPC: 11 cells, APPL1: 12 cells, Stx6: 12 cells, from at least 3 independent experiments). Values are means ±SEM. * *P* < 0.05, ** *P* < 0.01, *** *P* < 0.001.

### 3.5. Live imaging reveals that Par3 alters CNTNAP2 dynamics in intracellular compartments

Since Par3 appeared to localize CNTNAP2 to intracellular vesicles, we wanted to determine if CNTNAP2 is actively recruited by Par3 and if this association affects mobility. We used live cell imaging and fluorescence recovery after photobleaching (FRAP) to study these characteristics of CNTNAP2^+^/Par3^+^ and CNTNAP2^+^/Par3ΔPDZ_all_^+^ vesicles in COS7 cells. Consistent with our previous data (Figure 3), Par3 forms distinctively large punctate clusters when co-expressed with CNTNAP2 while Par3ΔPDZ_a_n retains a diffuse cytoplasmic staining (Figure 5a).

**Figure 5:**
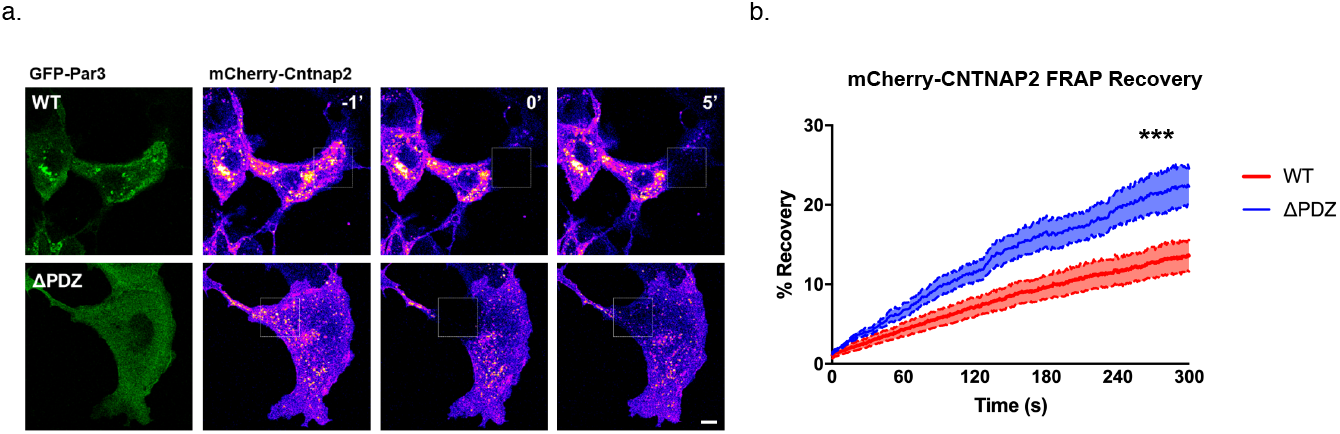
Par3 stabilizes CNTNAP2 within intracellular vesicles. (a) FRAP and time-lapse imaging (5 minutes) of live COS7 cells co-expressing mCherry-CNTNAP2 + GFP-Par3 (WT) or mCherry-CNTNAP2 + GFP-Par3ΔPDZ_all_ (ΔPDZ) (scale bar = 5 *μ*m). Green channel represents expression patterns of GFP-Par3 or GFP-Par3ΔPDZ_all_. Pseudo-color channel represents mCherry-CNTNAP2 before (−1’), during (0), or after (5’) FRAP bleaching in either condition. (b) Quantification of post-FRAP recovery in either condition by two-way ANOVA (WT: *n* = 8 cells, ΔPDZ: *n* =10 cells from 1 independent experiment). Values are means ±SEM. *** *P* < 0.001. WT τ=426.2s, ΔPDZ_all_ τ=361.3s

Time-lapse imaging of overexpressed mCherry-CNTNAP2 and GFP-Par3 in COS7 cells revealed a mixed population of large and immobile CNTNAP2^+^/Par3^+^ vesicles intermixed with smaller, more mobile ones (Supplemental Movie 1). Conversely, the immobile CNTNAP2^+^/Par3^+^ structures were abolished when GFP-Par3 was replaced with the Par3ΔPDZ_all_ mutant (Supplemental Movie 2). Using FRAP, we found CNTNAP2 mobility was increased when coexpressed with the Par3ΔPDZ_all_ mutant compared to WT. Five minutes following bleaching, mCherry-CNTNAP2 fluorescence recovered to twice the extent when coexpressed with GFP-Par3ΔPDZ_all_ compared to WT (Figure 5a-b). Additionally, the rate of recovery was nearly 20% faster with the Par3ΔPDZ_all_ mutant (WT: *τ* = 426.2s, APDZ_all_: *τ* = 361.3s). These data suggest that Par3 restricts CNTNAP2 mobility in a manner dependent on the PDZ domains of Par3.

### 3.6. Par3 affects the localization of CNTNAP2 in primary neuronal dendrites

Having shown that Par3 can influence CNTNAP2’s localization in COS7 cells, we set out to understand this relationship in cultured rat neurons. Because our previous study and others show CNTNAP2 to be more highly expressed in interneurons (Gao and Penzes, 2015; Mo et al., 2015; Karayannis et al., 2014), we focused our analysis on inhibitory neurons. We overexpressed Par3 in interneurons and studied its expression patterns as well as that of endogenous CNTNAP2. We confirmed the specificity of our interneuronal marker by co-staining with GABA (Supplementary Figure 2a). In line with our observations in cell lines, Par3 was largely diffuse in the cytoplasm but also formed large vesicular-like clusters (Figure 6a), consistent with endogenous staining patterns (Supplementary Figure 2b) (Zhang and Macara, 2006). On the other hand, overexpression of the Par3ΔPDZ_all_ mutant produced mostly a cytoplasmic pattern, suggesting the PDZ interaction is critical for clustering (Figure 6a). To ensure that our observations are robust, we confirmed the specificity of the CNTNAP2 antibody by staining *CNTNAP2*^−/−^ neurons (Supplementary Figure 3). Finally, as in COS7 cells, we occasionally found large aggregates of endogenous CNTNAP2 that clustered strongly with overexpressed Par3, but not Par3ΔPDZ_all_ (Figure 6a). Altogether, our data indicate that Par3 can direct localization of CNTNAP2 to punctate structures in neurons.

**Figure 6:**
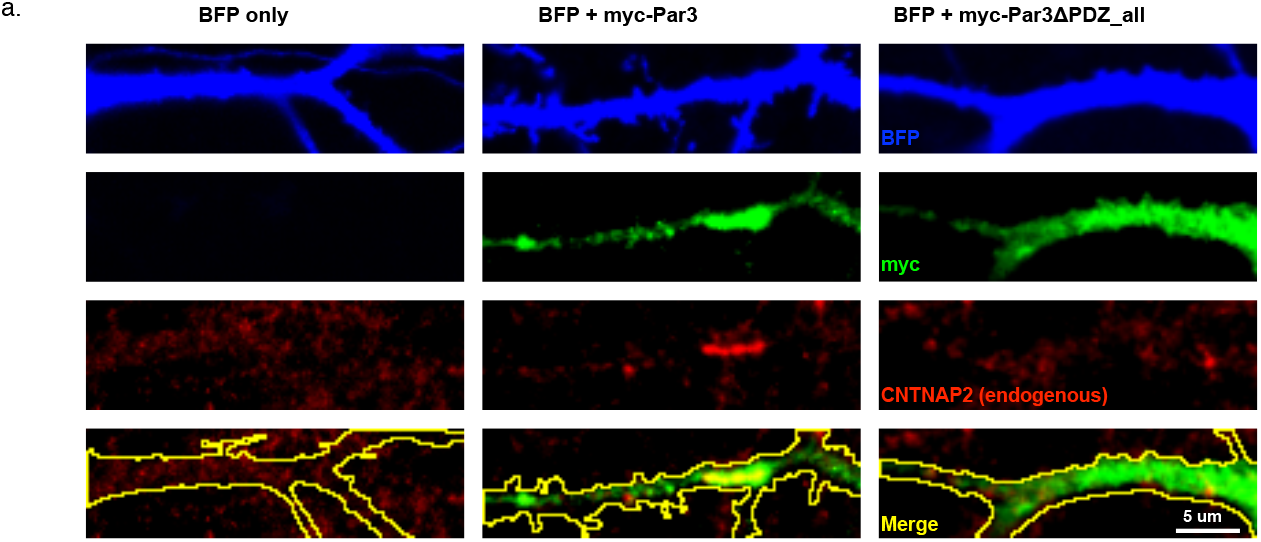
Overexpressed Par3 clusters endogenous CNTNAP2 in primary inhibitory neurons. (a) Representative images (from at least 3 independent experiments) of cultured primary inhibitory neurons transfected with blue fluorescent protein (BFP), alone or together with myc-Par3 or myc-Par3ΔPDZ_all_ (scale bar = 5 *μ*m).

## 4. Discussion

Par3 is part of a conserved signaling complex required for inducing polarity events critical for cellular function and survival, such as migration, apical-basal polarity establishment, and axon specification (Mertens et al., 2006). Par3, with its multi-domain scaffold partners, serves as a central recruiter of downstream signaling molecules in these processes. Our work expands on our knowledge of this function by characterizing the existence of a previously unknown interaction with CNTNAP2 identified by unbiased yeast two-hybrid screening.

CNTNAP2 has multi-faceted roles in neuronal migration, development, and maintenance; Par3 may serve as a spatial regulator for these functions (Zhang and Macara, 2006; Ahmed and Macara, 2017). In support of this model, we show that Par3 clusters CNTNAP2 into large vesicles via C-terminal PDZ binding in heterologous cells. These CNTNAP2+/Par3+ vesicles appear to be components of the clathrin-dependent endosomal trafficking pathway, within which Par3 stabilizes and spatially restricts CNTNAP2. Translating to neurons, it is possible that relative levels of Par3 at a local scale within a neuron may influence CNTNAP2’s function (Ruch et al., 2017). For example, Par3 within spines may influence local CNTNAP2 levels and function in that area (Zhang and Macara, 2006; Duman et al., 2013), leading to alterations in synaptic density, dynamics, or signaling (Gdalyahu et al., 2015; Varea et al., 2015). Additionally, we observed CNTNAP2 and Par3 colocalized structures in dendrites of inhibitory neurons (Figure 6). These structures were fairly sparse within single cells but also suggest an spatially regulated assembly model. Importantly, these colocalizing structures were absent when Par3 was replaced with the ΔPDZ_all_ mutant.

Mechanistically, we found multiple Par3 PDZ domains are necessary for proper CNTNAP2 binding. While this may be the result of artificial steric disruption, it is possible that such architecture exists to hold multiple CNTNAP2 molecules together within a tight spatial confine (Renschler et al., 2018). This could explaining the abnormally large size of CNTNAP2^+^/Par3^+^ vesicles upon overexpression and suggesting the possibility of a network assembly (Hayashi et al., 2009). This configuration may also be necessary to offset the naturally weak CNTNAP2-Par3 interaction, as seen by our fractionation results (Figure 2).

Yet, how exactly does Par3 mechanistically regulate CNTNAP2? The answer is difficult to determine since CNTNAP2 has multiple regions critical to trafficking. It has been established that threonine 1292 on CNTNAP2’s intracellular domain regulates endocytosis through protein kinase C (Bel et al., 2009) and that proper surface targeting of family member neurexin1 requires its PDZ-binding motif (Fairless et al., 2008). Furthermore, specific sites on CNTNAP2’s extracellular region are also critical for proper secretory and axonal trafficking (Pinatel et al., 2017; Falivelli et al., 2012). Our immunocytochemistry experiments link Par3-CNTNAP2 complexes with clathrin-coated endosomes, suggesting that Par3 may, via PDZ binding, be another component critical for CNTNAP2 internalization from the surface membrane. These questions will be addressed in future studies.

Both CNTNAP2 and Par3 play important roles in several steps of cortex development, including cell proliferation, migration, myelination, cell adhesion and synapse development (Bultje et al., 2009; Liu et al., 2018; Tep et al., 2012; Famulski et al., 2010; Varea et al., 2015; Penagarikano et al., 2011; Scott et al., 2017). While defects in all of these processes have been implicated in neurodevelopmental disorders, further work is needed to determine the specific processes where their interaction plays a key role. Future work should focus on the molecular pathways dependent on this newly identified CNTNAP2-Par3 interaction, and to elucidate the specific processes in which this partnership operates.

### Ethics Statement

All experiments were approved by the Institutional Animal Care and Use Committee of Northwestern University, protocol number IS0005992.

## Supporting information

Materials List

## Author Contributions

R.G. led the project. R.G. and C.P.P. performed all confocal imaging experiments. R.G., M.D.M., and M.P.F. performed biochemistry experiments. S.Y. performed live imaging experiments. P.P. supervised the project. R.G., C.P.P. and P.P. wrote the manuscript.

## Acknowledgments

This work was supported by grants NS100785 and MH097216 from the NIH-NIMH to P.P and F30MH096457 to R.G.

