## Supplementary material for "CNTNAP2 is targeted to endosomes by the polarity protein Par3": Materials List

| **Antibody** | **Species** | **Company, Cat# (RRID)** | **Dilution** | **Application** | **Figure** |
| --- | --- | --- | --- | --- | --- |
| FLAG | Mouse | Sigma; F1804  (AB_262044) | 1:1000 | WB | Fig 1c |
|  |  |  | 3 µg per sample | IP | Fig 1c |
| Myc | Mouse | Santa Cruz; sc-40  (AB_627268) | 1:100 | WB | Fig 1c, d |
|  |  |  | 3 µg per sample | IP | Fig 1d |
|  |  |  | 1:100 | ICC | Fig 3a  Fig 4a  Fig 6a  Sup Fig 2b, c |
| CNTNAP2 | Mouse | Neuromab; K67/25  (AB_10673031) | 3 µg per sample | IP | Fig 1b |
| CNTNAP2 | Rabbit | Millipore; AB5886-50UL  (AB_92118) | 1:400 | ICC | Fig 6a  Sup Fig 4 |
|  |  |  | 1:2000 | WB | Fig 1b, d |
| Par3 | Rabbit | Millipore; 07-330  (AB_2101325) | 1:1000 | WB | Fig 1b |
|  |  |  | 1:100 | ICC | Sup Fig 3b |
| IgG | Mouse | Santa Cruz; sc-2025  (AB_737182) | 3 µg per sample | IP | Fig 1b, c, d |
| GluA1 | Mouse | Millipore; mab2263  (AB_1977459) | 1:1000 | WB | Fig 2 |
| PSD95 | Mouse | Neuromab; K28/43  (AB_2292909) | 1:1000 | WB | Sup Fig 1b |
|  |  |  | 1:1000 | ICC | Sup Fig 3a |
| β-Tubulin | Rabbit | Developmental Studies Hybridoma Bank (DHSB); E7 (AB_528499) | 1:1000 | WB | Sup Fig 1b |
| dsRed | Rabbit | Clontech; 632496  (AB_10013483) | 1:1000 | ICC | Fig 3a |
| Clathrin Heavy Chain | Rabbit | Cell Signaling Technologies; #4796  (AB_10557412) | 1:1000 | ICC | Fig 3a |
| EEA1 | Rabbit | Cell Signaling Technologies; #3288  (AB_2096811) | 1:1000 | ICC | Fig 3a |
| Rab5 | Rabbit | Cell Signaling Technologies; #3547  (AB_2300649) | 1:1000 | ICC | Fig 3a |
| Caveolin-1 | Rabbit | Cell Signaling Technologies; #3267  (AB_2275453) | 1:1000 | ICC | Sup Fig 2b |
| APPL1 | Rabbit | Cell Signaling Technologies; #3858  (AB_2056989) | 1:1000 | ICC | Sup Fig 2c |
| Syntaxin-6 | Rabbit | Cell Signaling Technologies; #2869  (AB_2196500) | 1:1000 | ICC | Not shown |
| GOPC | Rabbit | Cell Signaling Technologies; #8576  (AB_10891809) | 1:1000 | ICC | Not shown |
| GFP | Chicken | Abcam (Cambridge, MA, USA); ab13970  (AB_300798) | 1:10,000 | ICC | Figure 6a |

| Primer | Sequence | Vector backbone | Notes |
| --- | --- | --- | --- |
| FLAG_CNTNAP2_1_F | 5’ CGCGGGCCCGGGATCCATGCAGGCGGCTCCGCGC 3’ | peGFP-N2 | FLAG-CNTNAP2; two amplicons fused with Infusion; BamHI and Not I |
| FLAG_CNTNAP2_1_R | 5 ‘CTTGTCGTCATCGTCTTTGTAGTCCGTCCAGGCTCT 3’ |  |  |
| FLAG_CNTNAP2_2_F | 5’ GACGATGACGACAAGGCTCCCTCCACGTCCCAAAA |  |  |
| FLAG_CNTNAP2_2_R | 5’ TCTAGAGTCGCGGCCGCTCAAATGAGCCATTCCTTTTTGCTTTCATCAATGGTCTC 3’ |  |  |
| FLAG_CNTNAP2_d4.1B_1_F | 5’ CGCGGGCCCGGGATCCATGCAGGCGGCTCCGCGC 3’ |  | FLAG-CNTNAPd4.1B; two amplicons fused with Infusion; BamHI and Not I |
| FLAG_CNTNAP2_d4.1B_1_R | 5’ GGCGGCGTCCGCGCTGAACATGTACCG GATCAGGAAGACCAGGGTGCACAGGATGGTGAAAAT 3’ |  |  |
| FLAG_CNTNAP2_d4.1B_2_F | 5’ AGCGCGGACGCCGCCATCATGAA 3’ |  |  |
| FLAG_CNTNAP2_d4.1B_2_R | 5’ TCTAGAGTCGCGGCCGCTCAAATGAGCCATTCCTTTTTGCTTTCATCAATGGTCTC 3’ |  |  |
| FLAG_CNTNAP2_dPDZ_F | 5’ CGCGGGCCCGGGATCCATGCAGGCGGCTCCGCGC 3’ |  | FLAG-CNTNAPdPDZ; BamHI and Not I |
| FLAG_CNTNAP2_dPDZ_R | 5’ TCTAGAGTCGCGGCCGCTCATTTGCTTTCATCAATGGTCTCTGTGAAGTTGGG 3’ |  |  |
| FLAG_CNTNAP2_dCT_F | 5’ CGCGGGCCCGGGATCCATGCAGGCGGCTCCGCGC 3’ |  | FLAG-CNTNAPdCT; BamHI and Not I |
| FLAG_CNTNAP2_dCT_R | 5’ TCTAGAGTCGCGGCCGCTCAGATCAGGAAGACCAGGGTGCACAGGAT 3’ |  |  |
| mCherry_CNTNAP2_1_F | 5’ CGCGGGCCCGGGATCCATGCAGGCGGCTCCGCGC 3’ |  | mCherry-CNTNAP2; two amplicons fused with Infusion; BamHI and Not I |
| mCherry_CNTNAP2_1_R | 5’ CTCGCCCTTGCTCACCGTCCAGGCTCTGCAGAGGCA 3’ |  |  |
| mCherry_CNTNAP2_2_F | 5’ GTGAGCAAGGGCGAGGAGGATAACATGGC 3’ |  |  |
| mCherry_CNTNAP2_2_R | 5’ TCTAGAGTCGCGGCCGCTCAAATGAGCCATTCCTTTTTGCTTTCATCAATGGTCTC 3’ |  |  |
| Myc_PAR3_FL_F | 5’ CGCGGGCCCGGGATCCATGGAACAGAAACTCATCTCTGAAGAGGATCTGT 3’ |  | Myc-Par3 full-length; fused with infusion; BamHI and Not I |
| Myc_PAR3_FL_R | 5’ TCTAGAGTCGCGGCCGCTCAGGAATAGAAGGGCCTCCCTTTCT 3’ |  |  |
| Myc_PAR3_PDZ1_1_F | 5’ CGCGGGCCCGGGATCCATGGAACAGAAACTCATCTCTGAAGAGGATCTGT 3’ |  | Myc-Par3ΔPDZ1; two amplicons fused with Infusion; BamHI and Not I |
| Myc_PAR3_PDZ1_1_R | 5’ ATCATCCAGAGAAAAGTTGGGTATATGCTCCA 3’ |  |  |
| Myc_PAR3_PDZ1_2_F | 5’ TTTTCTCTGGATGATGCAGCAAATAAAGAGCAGTATGAACAACTA 3’ |  |  |
| Myc_PAR3_PDZ1_2_R | 5’ TCTAGAGTCGCGGCCGCTCAGGAATAGAAGGGCCTCCCTTTCT 3’ |  |  |
| Myc_PAR3_PDZ2_1_F | 5’ CGCGGGCCCGGGATCCATGGAACAGAAACTCATCTCTGAAGAGGATCTGT 3’ |  | Myc-Par3ΔPDZ2; two amplicons fused with Infusion; BamHI and Not I |
| Myc_PAR3_PDZ2_1_R | 5’ AAGCCTCTTGCCTATTTTTTTGGTGTTAT 3’ |  |  |
| Myc_PAR3_PDZ2_2_F | 5’ ATAGGCAAGAGGCTTCAGGAAGACGCCTTCCACCCA 3’ |  |  |
| Myc_PAR3_PDZ2_2_R | 5’ TCTAGAGTCGCGGCCGCTCAGGAATAGAAGGGCCTCCCTTTCT 3’ |  |  |
| Myc_PAR3_PDZ3_1_F | 5’ CGCGGGCCCGGGATCCATGGAACAGAAACTCATCTCTGAAGAGGATCTGT 3’ |  | Myc-Par3ΔPDZ3; two amplicons fused with Infusion; BamHI and Not I |
| Myc_PAR3_PDZ3_1_R | 5’ AAATGTCAGAAATTCCCTGGTGCCATCAGG 3’ |  |  |
| Myc_PAR3_PDZ3_2_F | 5’ GAATTTCTGACATTTCTTATTGTTGCAAGGAGAATAAGCAAGT 3’ |  |  |
| Myc_PAR3_PDZ3_2_R | 5’ TCTAGAGTCGCGGCCGCTCAGGAATAGAAGGGCCTCCCTTTCT 3’ |  |  |
| Myc_PAR3_PDZall_1_F | 5’ CGCGGGCCCGGGATCCATGGAACAGAAACTCATCTCTGAAGAGGATCTGT 3’ |  | Myc-Par3ΔPDZ_all_; two amplicons fused with Infusion; BamHI and Not I |
| Myc_PAR3_PDZall_1_R | 5’ ATCATCCAGAGAAAAGTTGGGTATATGCTCCA 3’ |  |  |
| Myc_PAR3_PDZall_2_F | 5’ TTTTCTCTGGATGATCTTATTGTTGCAAGGAGAATAAGCAAGT 3’ |  |  |
| Myc_PAR3_PDZall_2_R | 5’ TCTAGAGTCGCGGCCGCTCAGGAATAGAAGGGCCTCCCTTTCT 3’ |  |  |
| GFP_PAR3_F | 5’ TACCGGACTCAGATCTATGGTGAGCAAGGGCGAGGAG 3’ |  | GFP was inserted via Infusion into BglII and BamHI on Myc-Par3 (see above) backbone |
| GFP_PAR3_R | 5’ TCTGTTCCATGGATCCCTTGTACAGCTCGTCCATGCCGAGAG 3’ |  |  |
