## Supplementary Information for "CNTNAP2 is targeted to endosomes by the polarity protein Par3"

**Supplementary Methods**

*Primary neuronal culture from postnatal mouse.* For supplementary figure 4, WT and *Cntnap2^-/-^* neurons were used. Briefly, P0-1 mouse pups were decapitated and brains dissected on ice in cold Liebovitz’s L-15 media containing penicillin/streptomycin. After removal of meninges, cortices were separated from each hemisphere and placed into a tube containing L-15. Tissue was digested using 0.25% trypsin for 10 minutes at 37°C. Trypsin was removed and replaced with 1 mL plating media (DMEM containing 10% fetal bovine serum, 1x GlutaMAX (Gibco), 0.6% D-glucose (w/v), penicillin/streptomycin) Cortices were dissociated by 10 strokes using a P1000 pipette. Plating media was added to a volume of 10 mL and passed through a cell strainer. Cells were counted and seeded on 18mm poly-D-lysine coated #1.5 coverslips.

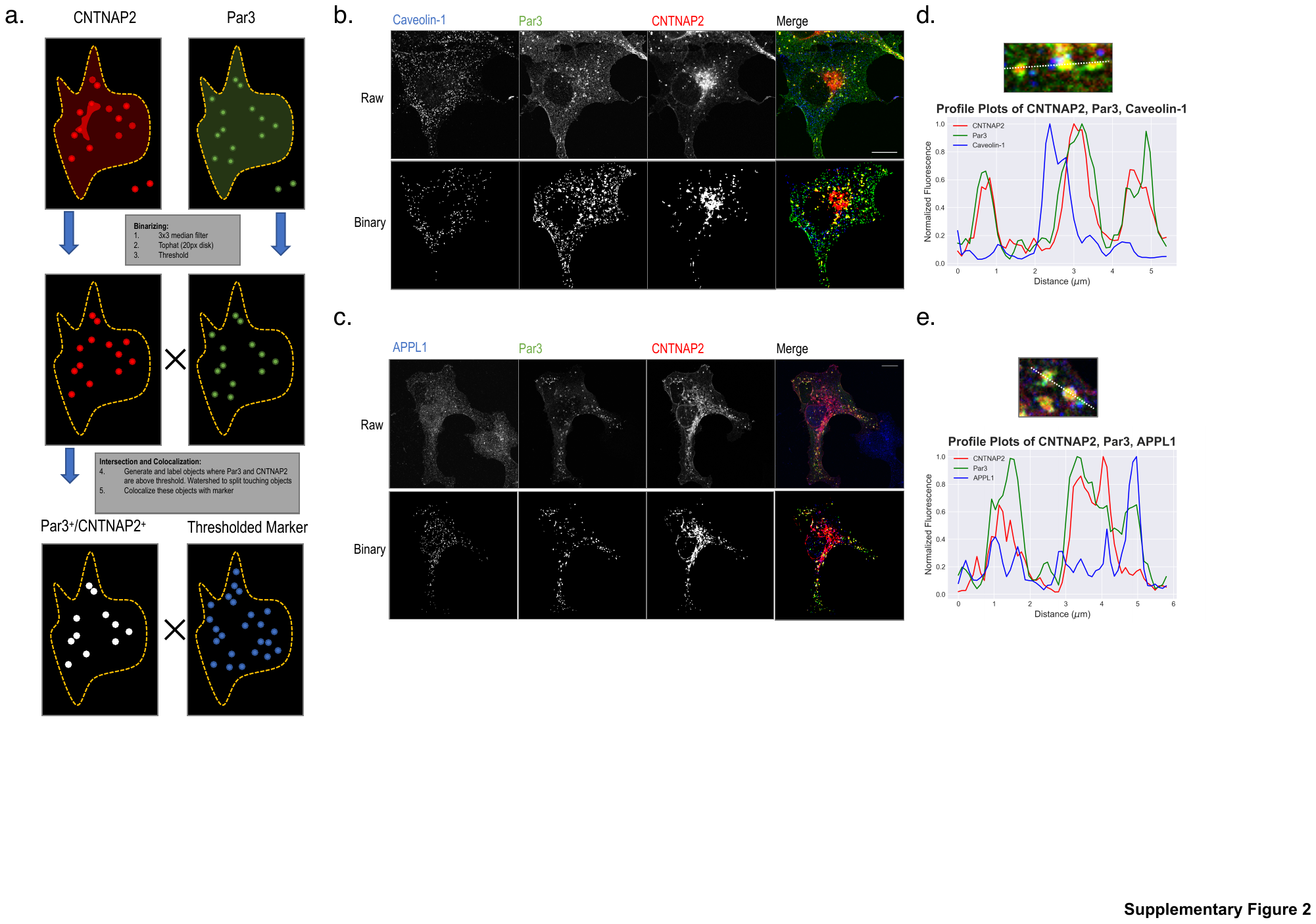
**Supplementary Figure 1.** Imaging workflow and examples of low co-localizing markers. (a) Image processing workflow to analyze colocalization: ROIs were drawn encompassing whole cells on single optical sections using BFP cell-fill as a guide. Images were processed using a 3x3 median filter and 20-pixel disk tophat transformation. Thresholding was performed on each individual ROI using the Triangle thresholding method or Otsu’s method (<5% of images). Colocalization was then measured using the resulting binary images. (b-c) Representative images (top rows) and their resulting binaries (bottom rows) are shown for Caveolin-1 (b) and APPL1 (c). (d-e) Profile plots for Caveolin-1 (d) and APPL1 (e) with magnified images (top) with dotted lines indicating line scans and the measured profile plots (bottom). Scale bars = 15 µm.

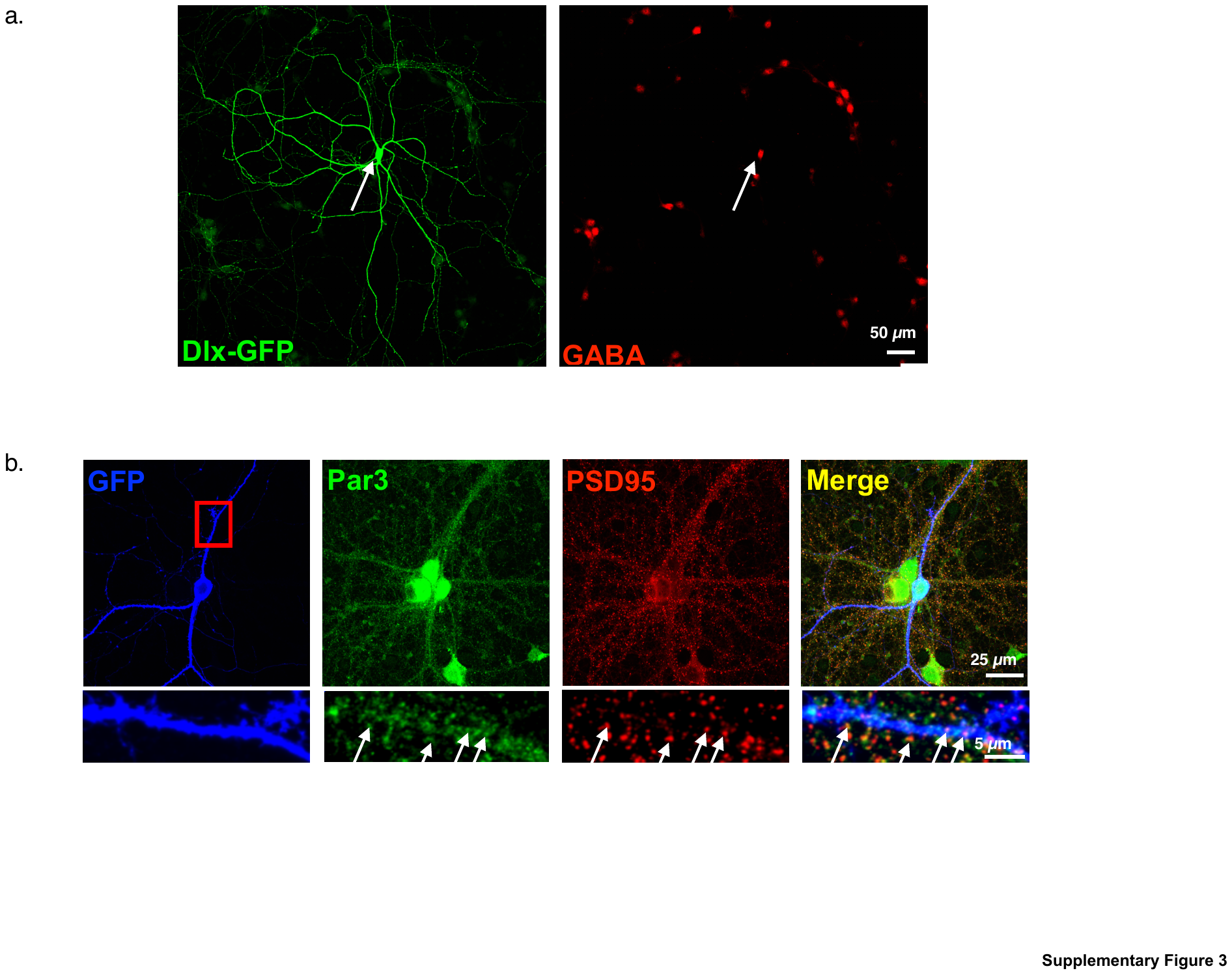

**Supplementary Figure 2.** Par3 is expressed in GABAergic inhibitory neurons (a) Interneuron-specific Dlx driven GFP shows expression in neurons positively stained for GABA. (b) Endogenous Par3 immunostaining colocalizes with PSD95 (white arrows) in GFP labeled interneurons.

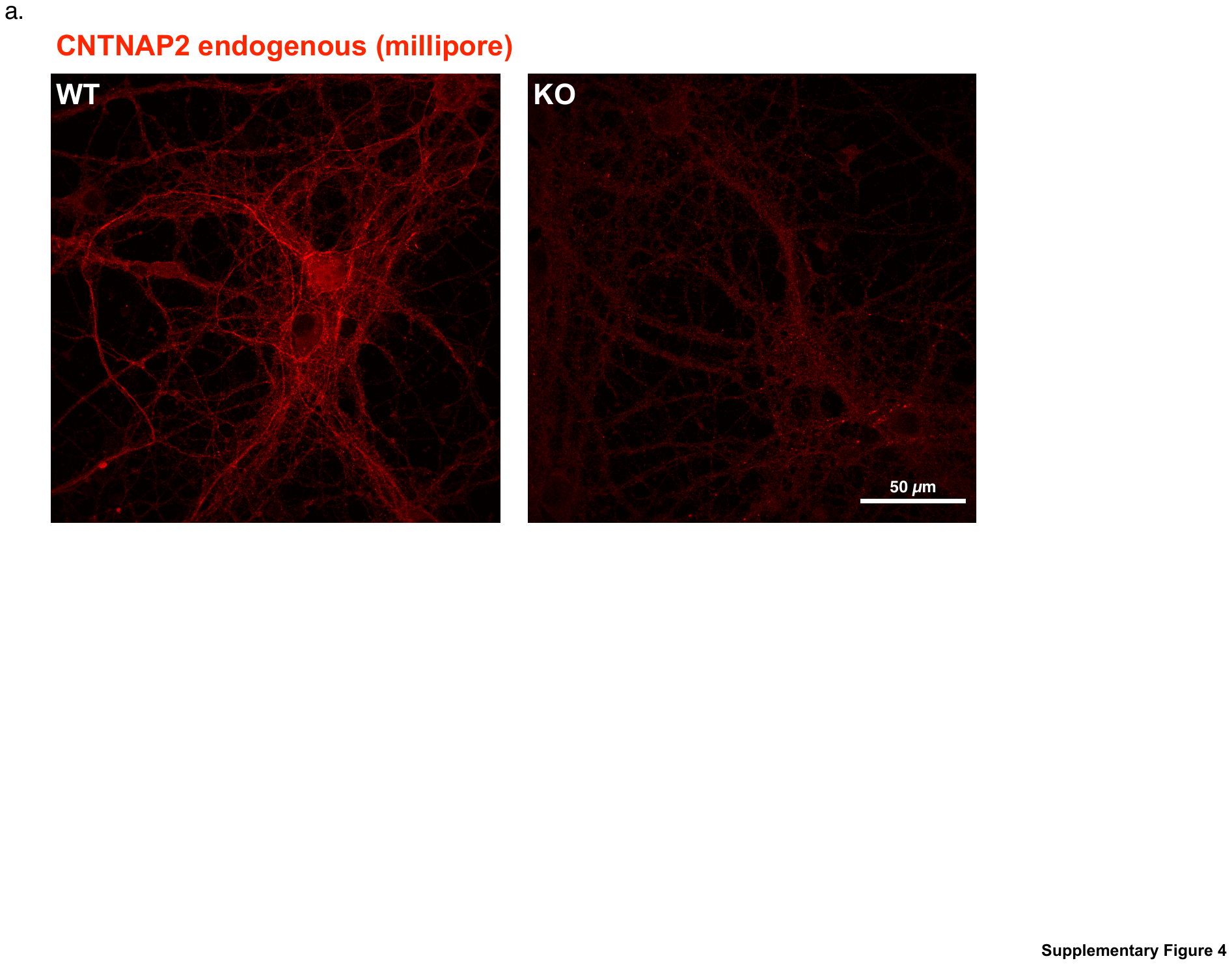

**Supplementary Figure 3.** CNTNAP2 antibody is specific for neuronal CNTNAP2. (a) Immunofluorescent staining of CNTNAP2 in wild type and *CNTNAP2* knockout mouse neurons.

**Supplemental Movie 1.** Representative live pre-bleach recording of a COS7 cell overexpressing mCherry-CNTNAP2 (magenta) and GFP-Par3 (green).

**Supplemental Movie 2.** Representative live pre-bleach recording of a COS7 cell overexpressing mCherry-CNTNAP2 (magenta) and GFP-Par3ΔPDZ_all_ (green).
